# Lipogenic Mothers Alter Offspring Gut Microbiome Development via Breastmilk-Mediated Microbial Transfer

**DOI:** 10.1101/2025.01.21.634074

**Authors:** Jianhong Cao, Abdulrahim Umar, Zheng Yu

## Abstract

Mammals provide breastmilk as the primary source of nutrition for their offspring, which plays a crucial role in the development of gut microbiota (GM) and the establishment of immunity. Therefore, maternal dietary conditions during lactation significantly influence GM development. This study investigates the impact of a maternal high-fat diet on GM development in their offspring. Tissue and fecal samples were collected from both pups and their mothers (dams) that were fed a high-fat and a controlled diet. Their amplicon sequencing variants were used to perform comprehensive bioinformatic analysis.

Our findings revealed breastmilk’s protective effect, inhibiting the proliferation of pathogenic bacteria, and regulating gut homeostasis during lactation. However, lipogenic dams induce microbial dysbiosis in their pups leading to an increased relative abundance of gut pathogenic bacteria leading to a heightened risk of chronic diseases. In contrast, pups from dams on a controlled diet exhibited an improved relative abundance of beneficial bacteria, including *Muribaculaceae*, *Faecalibaculum*, and *Bifidobacterium*. However, GM transferred from lipogenic dams adversely affected pups’ GM development and stability, leading to several physical consequences.

This research revealed valuable insights into the relationship between breastmilk and gut microbiota, as specific genera were highly expressed at the beginning of lactation. This highlights the importance of timely interventions for the development of offspring gut microbiota to enhance overall gut health.

## 1.0 Introduction

The human gut microbiome and its role in both health and disease have been the subject of extensive research over the years, establishing its involvement in human metabolism, nutrition, physiology, and immune function [1]. The development of the gut microbiome is influenced by various factors, including diet and early-life exposures. Maternal-offspring exchanges of microbiota during breastfeeding facilitate the transfer of gut-associated obligate anaerobic genera, such as *Bifidobacterium*, *Bacteroides*, *Parabacteroides*, and members of the Clostridia family (including *Blautia*, *Clostridium*, *Collinsella*, and *Veillonella*), observed in maternal feces, breastmilk, and neonatal feces [2,3].

*Enterococcus* and *Bifidobacteria* which are microorganism that breakdown human milk oligosaccharides, make up the majority of an infant’s gut microbiome when they are exclusively breastfed or fed formula. As a result, early GM formation is greatly influenced by neonates feeding [4,5].

Maternal consumption of a diet high in saturated fatty acids (SFA) and low in polyunsaturated fatty acids (PUFA) during pregnancy can adversely affect placental function, contributing to the development of placental and fetal metabolic dysfunction, regardless of maternal body composition [6]. However, the breastmilk of individuals consuming a healthy and balanced diet fosters the proliferation of beneficial bacteria. Therefore, high-fat dietary habits during lactation may predispose offspring to microbial dysbiosis leading to renal and metabolic injuries later in life [7].

The GM engages in a bidirectional interaction with the host immune system, playing a crucial role in promoting the maturation of the host’s immune response [8,9]. Notably, a maternal high-fat diet (MHFD) significantly alters the GM, impacting its composition, diversity, and richness. These changes can influence the infant’s gut microbiome acquisition and increase the risk of developing metabolic disorders, defective genes, and reproductive health [10]. Moreover, during pregnancy the influence of high-fat diets may lead to modifications in embryonic brain metabolites and affect adolescent behavior [11,12]. Furthermore, MHFD can lead to gut microbial dysbiosis, which contributes to an elevated inflammatory environment during pregnancy and lactation. This disruption can have a significant effect on the neurodevelopment of the offspring, impacting both prenatal and postnatal stages of the offspring [13].

In an animal study, MHFD was reported to be a significant risk factor for systemic inflammation and hepatic steatosis (fatty liver disease) in offspring mice [14]. This dietary risk factor adversely affects lipid metabolism in the offspring, potentially leading to elevated lipid levels, fat accumulation in various organs, and increased systemic inflammation. Unfortunately, maternal exposure to high-fat diets, diabetes, and obesity in the environment can have lasting effects on offspring [15]. In a cellular context, Wei et al. found that offspring may be more susceptible to cellular DNA damage due to MHFD intake [16].

Considering these challenges, we investigated the impact of maternal lipogenic diet-induced obesity on the offspring’s gut microbiome development, with a focus on the role of breastmilk-mediated microbial transfer. We hypothesized that high-fat dams, characterized by an imbalanced lipid profile, would transfer a distinct microbial community to their offspring via breastmilk, ultimately shaping the developing gut microbiome.

By employing 16S rRNA gene sequencing, we aim to elucidate the specific taxonomic alterations in the offspring’s gut microbiome due to MHFD exposure. This study provides valuable insights into the complex interplay between maternal nutritional status, breastmilk composition, and the establishment of the offspring’s gut microbiome, with implications for understanding the early-life origins of metabolic disorders.

## 2.0 Methods

### 2.1 Animals experiment

The experiment described in this manuscript was performed in accordance with the guidelines outlined in the Guide for the Care and Use of Laboratory Animals and was approved by the Institutional Ethics Committee for Animal Procedures of Central South University (CSU-2024-0315). The C57BL/6 mice were purchased from Hunan Sleek Jingda (SLAC), Changsha, China, and the mice were housed separately according to their groups in individually ventilated cages (IVCs) with free access to food and water for the potential delivery of offspring. All mice were raised in plastic cages covered with metal fences in the light (12/12 h light/dark cycle), humidity (60-70%), and temperature-controlled (25 ± 5 °C) facility under specific pathogen-free conditions.

The group consuming the high-fat diet was designated as HFD and the controlled diet was designated as CD. About 6 weeks old 8 female mice (HFD: n=4, CD: n=4) were bred and their subsequent pups were allowed to age until either day 0 (HFD: n=3, CD: n=3), 2 weeks (HFD: n=3, CD: n=3). Dams were randomly euthanized at age 18 weeks at random while subsequent pups were euthanized at each timepoint chosen at random, and samples were collected immediately after euthanasia. An additional 9 pup fecal samples (HFD: n=4, CD: n=5) were also collected at 3, 4, and 5 weeks of age from colony pups for comparison to the dam-pup matched samples.

### 2.2 Sample collection

All maternal fecal samples were collected at intervals within the periods (adaptation, pre-pregnancy, pregnancy, and post-pregnancy), the two most distal fecal pellets in the rectum were collected and placed in a 2 ml sterile EP tubes appropriate for homogenization of the sample. Pup feces were collected as described for dams at weeks 3, 4, and 5 of age. Pup proximal small intestine and ileal samples were collected at day0 and week 2 of age by excising 2 cm of the ileum proximal to the ileocecal junction and collecting into a 2 ml sterile EP tubes due to the small size of the ileal lumen. All samples were placed on ice immediately following collection and samples were stored in a −80 °C freezer until DNA was extracted.

### 2.3 DNA extraction

All sample tissue DNA was extracted using OMEGA Soil DNA Kit (M5636-02) (Omega Bio-Tek, Norcross, GA, USA) according to the manufacturer’s protocol and stored at −20°C prior to further analysis. The quantity and quality of DNA yield were measured by NanoDrop NC2000 spectrophotometer (Thermo Fisher Scientific, Waltham, MA, USA) and agarose gel electrophoresis respectively. Then normalized to a consistent concentration and volume prior to submission for downstream processing.

### 2.4 16S rRNA Library Preparation and Sequencing

Bacterial 16S rRNA amplicons were constructed via polymerase chain reaction (PCR) amplification of the V3-V4 region of the 16S rRNA gene with the universal primer set 338F (5’-ACTCCTACGGGAGGCAGCA-3’) and reverse primers 806R (5’-GGACTACHVGGGTWTCTAAT-3’) and flanked by Illumina standard adapter sequences as in a method previously described elsewhere (Walters et al., 2011). Dual-indexed forward and reverse primers were used in all sample reactions. The qualified libraries were then sequenced on the Illumina NovaSeq platform with the NovaSeq-PE250 sequencing strategy at Shanghai Personal Biotechnology Co., Ltd. (Shanghai, China). After sequencing, the data was decomposed into appropriate samples based on barcodes, and the sequences were imported into downstream software.

### 2.5 Data Analysis and Graphing

Quality control and denoising of raw reads were performed based on the standard amplicon pipeline as described previously in QIIME2 [18]. The microbiome analysis output (feature table and taxonomy annotation table) was used for further data analysis. To calculate significant changes in alpha diversity and relative abundance (RA) of maternal and offspring samples, we utilized ANOVA on ranks within each diet due to a lack of normality. Bacteria genera with significant changes in relative abundance were discovered by Linear Discriminant Analysis (LDA) Effect Size (LEfSe) (http://huttenhower.sph.harvard.edu/galaxy/) [19]. The threshold of significance was set at 0.05, while the threshold of the LDA score was set at 4.0. The analysis of Spearman correlation for genera with significant changes in relative abundance was calculated and visualized. Community richness, the number of unique amplicon sequence variants (ASVs) within a sample, using PAST software [20]. An ANOVA on ranks was performed using SigmaPlot 14.0 with a Dunn’s post hoc analysis for pairwise comparisons. Analysis of similarity (ANOSIM) and One-way permutational multivariate analysis of variance (PERMANOVA) was used to test for significant differences in beta diversity of samples and provide pairwise comparisons of beta-diversity. PERMANOVA testing was performed using PAST software using Bray-Curtis’s similarities. The Bray-Curtis similarity of pup fecal samples to maternal samples were tested for significant differences within each diet using two-way ANOVAs for the age using SigmaPlot 14.0. The threshold of the P-value corrected by the Benjamin and Hochberg false discovery rate (FDR) was 0.05.

## 3.0 Results

### 3.1 Experimental overview, physical and morphological changes in dams and their offspring

We successfully executed the experimental plan according to (Fig. 1A). Dams on the HFD exhibited elevated fasting blood sugar throughout the breeding period, with significant increases noted during the 7^th^ and 11^th^ weeks, corresponding to pregnancy and lactation, respectively (Fig. 1C). In contrast, CD dams experienced greater weight gain during pregnancy and lactation compared to their HFD counterparts (Fig. 1B).

**Figure 1:**
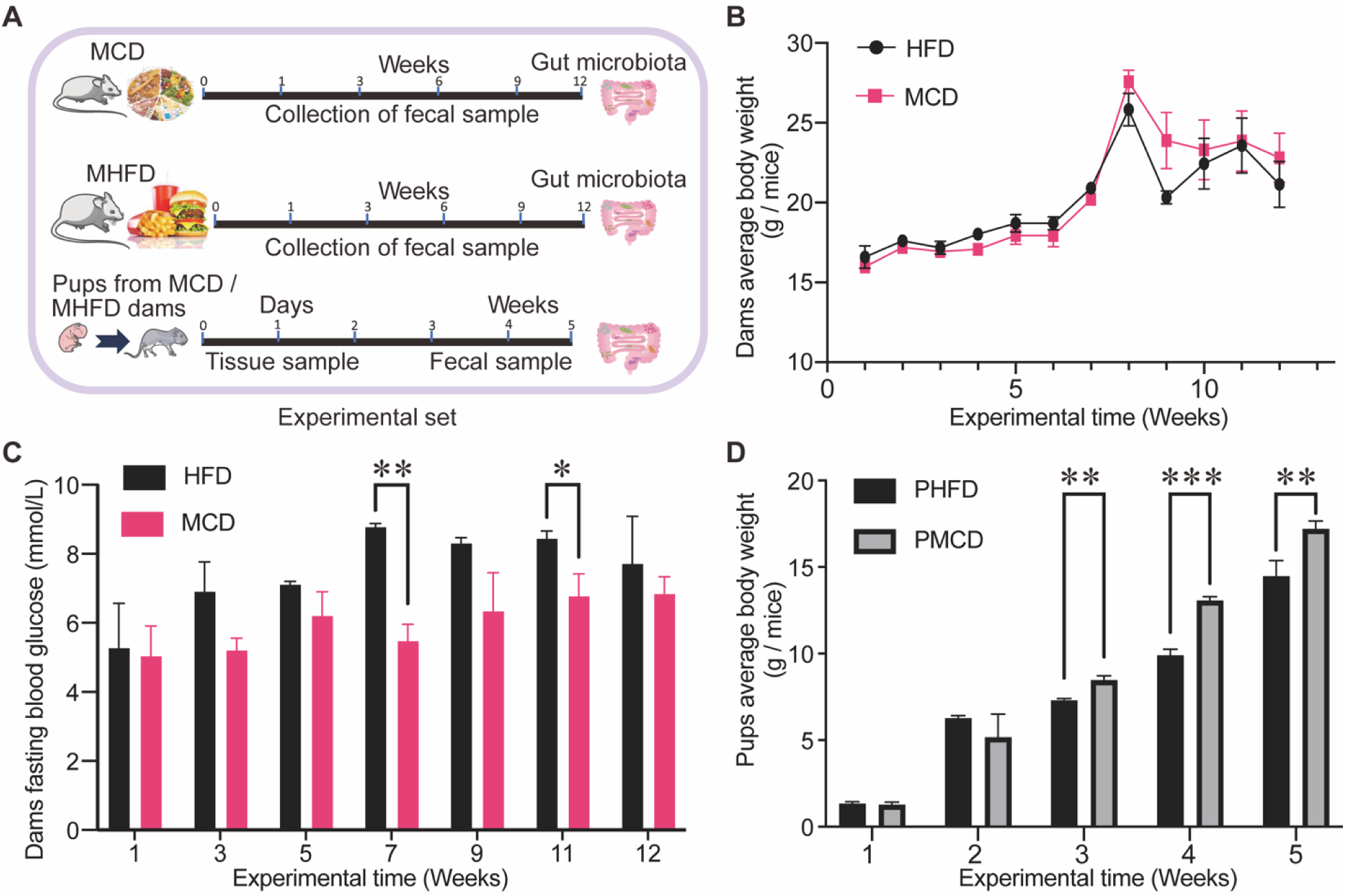
HFD induced significant changes in characteristic features observed in dams and pups. (A) Experimental set-up scheduled for the study. (B) Changes in relative body weight of the dams among the two groups. (C) Changes in fasting blood glucose level of dams among the two groups. (D) Body weight changes in two groups of offspring mice. Using multiple t-test comparison, *** indicates P < 0.001, ** indicates P<0.01, and * indicates P < 0.05.

There was no significant difference in the body weight of the pups at birth; however, significant differences in the body weight emerged during lactation and throughout the development periods (Fig. 1D). This observation underscores the impact of HFD on the developmental processes of the offspring.

### 3.2 Lipogenic dams induce microbial-level compositional changes during offspring GM Development

Fecal analysis of offspring from MHFD and MCD were changing in phylum composition on day0 and week2. Even though, the microbial composition was relatively stable from week3, the relative abundance of each phylum and genera differed from each other (Fig. 2A, B). Notably, *Bacteroidota* did not exist before lactation and began to colonize the offspring’s gut after lactation (Fig. 2A, C). In contrast, *Acinetobacter*, present in dams with MHFD, was transferred to pups prior to lactation (Fig. 2B); however, its RA diminished completely as the pups continued to develop. In a specific context, the microbial genera *Bacteroides*, *Parabacteroides*, and *Muribaculaceae* exhibited a higher RA compared to *Lactobacillus*, which showed high RA only in week2 and *Atopobiaceae*, which showed significantly lower RA at each timepoint during the development. Although, *Bacteroides*, *Muribaculaceae*, and *Atopobiaceae* maintained stable RA between the 4^th^ and 5^th^ weeks (Fig. 2B).

**Figure 2:**
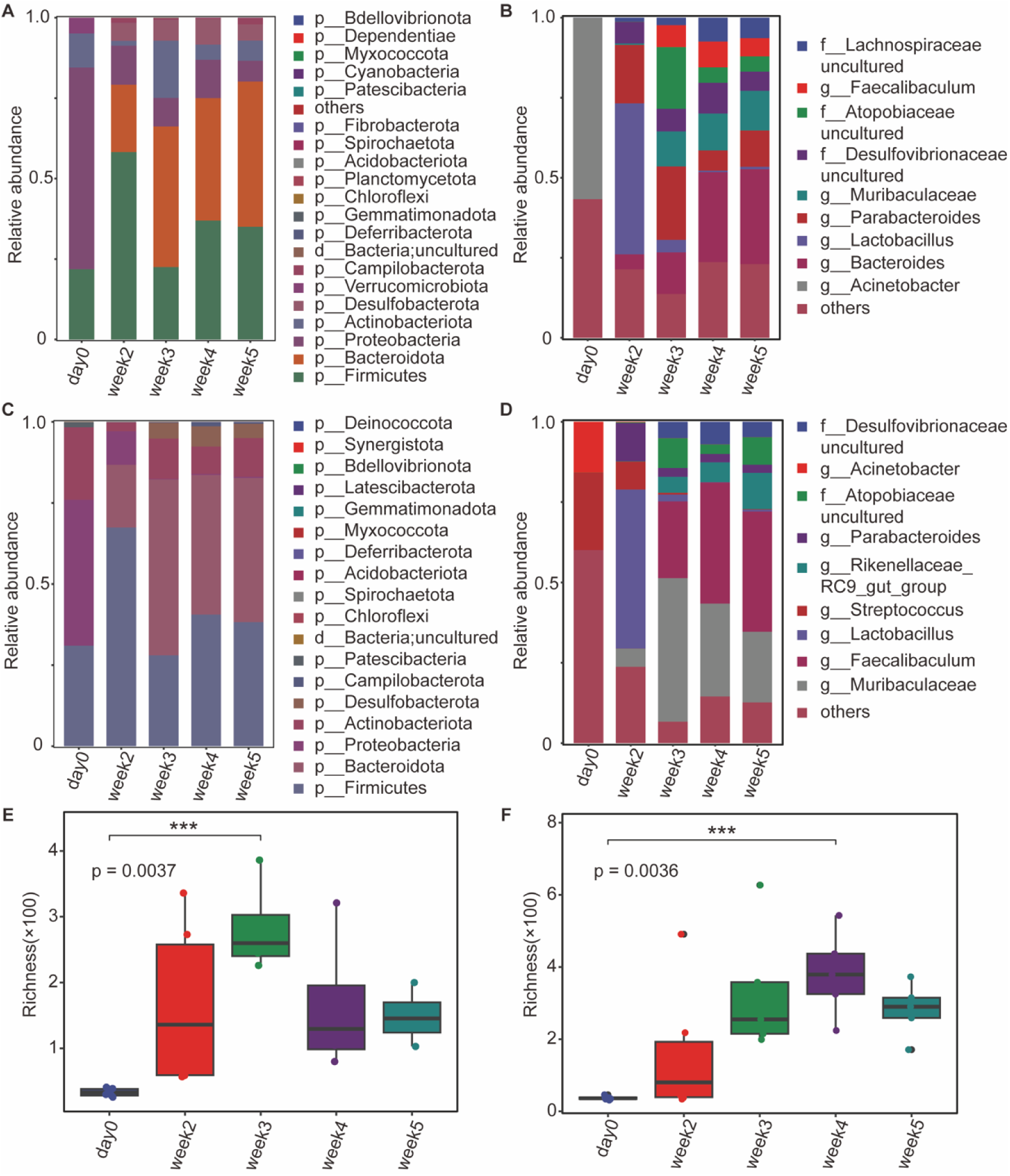
Dams high-fat diet induced microbial dysbiosis in pups. Changes in relative abundance (RA) of phyla (A) and genera (B) in pups from MHFD. Changes in RA of phylum (C) and genera (D) in pups from PMCD. The X-axis represented the experimental days. The Y-axis represented the relative abundance of different phyla and genera. Modules in different colours separated different phyla or genera. “others” represented uncultured phyla or genera yet to be fully classified. Timepoint richness alpha diversity represented pups from PHFD (E) and pups from PMCD (F) respectively and error bars represent the standard error of mean (SEM).

Offspring from the MCD and MHFD displayed a stable RA of *Bacteroidota*, *Actinobacteriota*, and *Proteobacteria* (Fig. 2C) from week3 of the development timepoints. Additionally, a higher RA of *Muribaculaceae* and *Faecalibaculum* was observed (Fig. 2D) when compared to its genus-level composition (Fig. 2B). Unique RAs of *Streptococcus* and *Rikenellaceae_gut_group* were also noted at each developmental stage (Fig. 2D).

The GM derived from amniotic fluid (pups’ residence during pregnancy) and those derived from breastmilk may have significantly altered the GM dynamics of the pups by contributing *Firmicutes* and *Bacteroidota*, while *Proteobacteria* transferred via amniotic fluid were completely absent in both groups as the development continued to progress.

These phylum- and genus-level compositional changes were verified using Principal Coordinates Analysis (PCoA), which demonstrated significant variations between the treatments (HFD and CD) administered to the dams. Alpha diversity analysis indicated marked differences in microbial communities between the two dietary groups (Supplementary Fig S1).

An ANOVA on ranks test was conducted to assess differences in microbial richness at each developmental stage, defined as the number of unique ASVs within a sample. As expected, the richness of pups’ fecal samples increased exponentially from day0 to the 3rd week after weaning. However, the pups from the maternal high-fat diet (PHFD) group exhibited a rapid decline in microbial richness, while the pups from the maternal control diet (PMCD) group continued to increase their microbial richness despite dams weaning (Fig. 2E, F). The results for PMCD indicated higher microbial richness with significant differences observed between day_0 and the 4^th^ week (Fig. 2F), in contrast to the PHFD group, which showed a lower microbial richness with significant differences by the 3^rd^ week and a rapid decline following dams weaning (Fig. 2E).

### 3.3 Offspring from MHFD Express High-level Relative Abundance of Pathogenic Genera

The cluster heat map of pups’ development time points significantly distinguishes the RA of GM between PHFD and PMCD. Fecal analysis revealed a predominance of beneficial genera including *Muribaculaceae*, *Faecalibaculum*, *Bifidobacterium*, *Alloprevotella*, *Alistipes*, and *Odoribacter* in the PMCD group sharing a common phylogenetic origin. In contrast, the PHFD group exhibited higher RA of mostly pathogenic genera including *Escherichia-Shigella*, *Enterococcus*, *Blautia*, *Citrobacter*, *Bacteroides*, *Clostridium* (innoculum_group), *Parasutterella*, *Erysipelatoclostridium*, *Desulfovibrionaceae*, and *Enterobacteriaceae*. This expression of different phylogenetic bacteria genera was more pronounced during the 3^rd^ to 5^th^ weeks of their pups’ development (Fig. 3A). This study traced the RA of these genera in pups to have been vertically transferred via breast milk from their mothers. Notably, *Faecalibaculum* and *Bifidobacterium* were more prevalent in dams fed a controlled diet, while *Escherichia-Shigella*, *Blautia*, and *Erysipelatoclostridium* were more abundant in dams consuming a HFD (Supplementary FS2). It is evident that the expressions of this genus were not observed before lactation; thus, microbiota transfer during lactation played a significant role in the initiation and development of pup’s gut microbiota (Fig. 3A).

**Figure 3:**
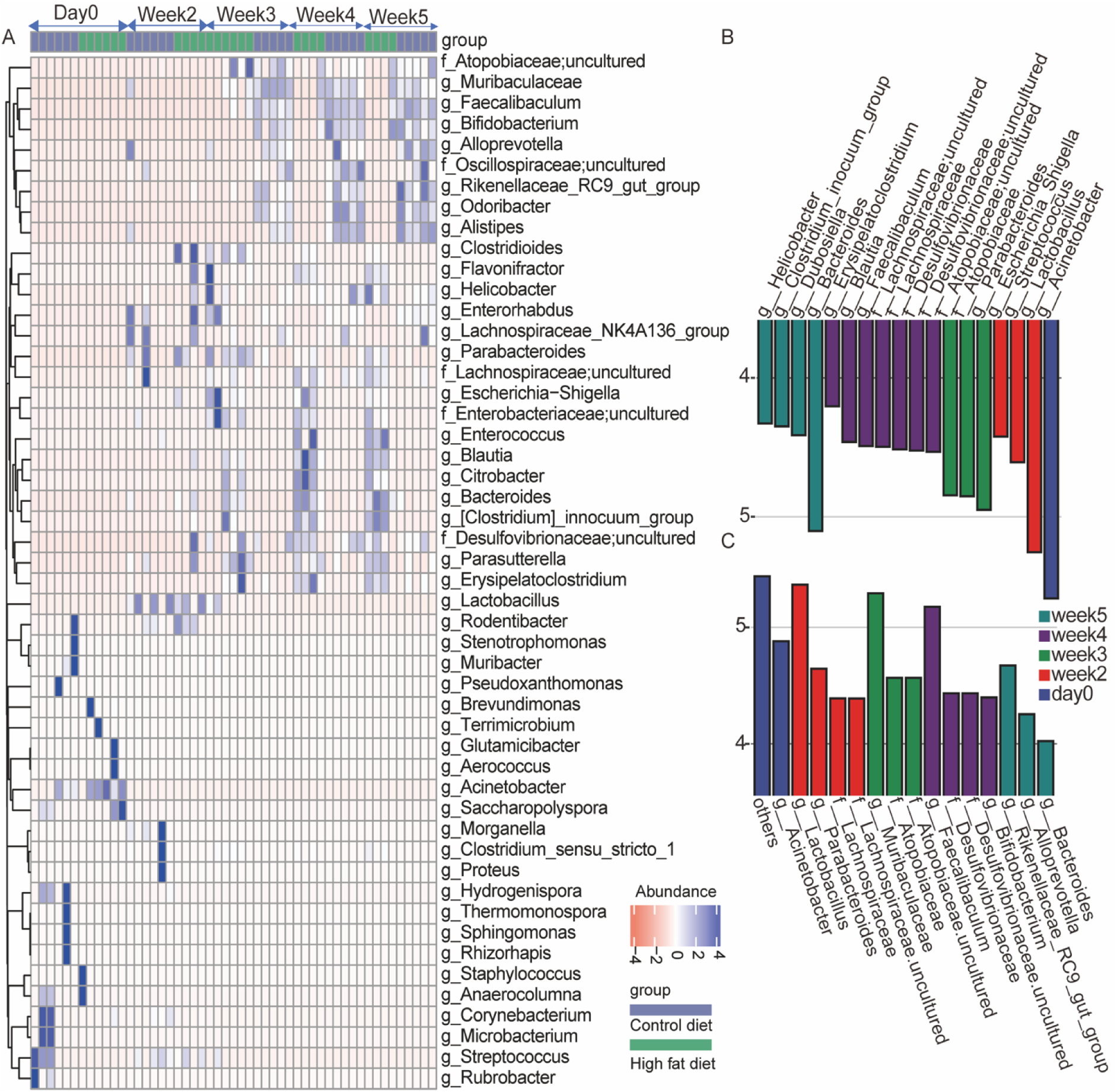
Pups’ genera with significant differences in stages of development. (A) cluster heat map showing the abundance level of genera from different and similar taxonomic groups. Different colours indicated significant changes in the relative abundance of different groups. The blue colours represent a highly significant change in corresponding genera. A histogram with linear discriminant analysis (LDA) scores in five groups for pups from MHFD (B) and PMCD (C) respectively. Taxa highlighted in different colours indicated overrepresentation in the corresponding groups. The threshold of significance was set at 0.05. The threshold of the LDA score was set at 4.

Other unique expressions included *Lactobacillus*, which was highly expressed in both PHFD and PMCD groups during the 2^nd^ week of GM development. Additionally, *Flavonifractor* and *Helicobacter* exhibited high RA in the PHFD group during the 2^nd^ week, while *Streptococcus*, *Acinetobacter* were predominantly expressed at day0 in the pups’ GM (Fig. 3A).

LEfSe analysis provided a detailed description of the relatively abundant genera at each time point of the pups’ development. Prior to lactation, both PHFD and PMCD contained the genus *Acinetobacter* at day0 (Fig. 3B, C). Following the onset of lactation, the GM of pups from both the PMCD and PHFD groups harbored *Lactobacillus* during the 2^nd^ week. However, pathogenic bacteria genera such as *Streptococcus* and *Escherichia-Shigella* were also relatively dominant in the 2^nd^ week. The LEfSe analysis for the MHFD group also highlighted the abundance of *Escherichia-Shigella* and *Lachnospiraceae* during pregnancy and lactation periods respectively, whereas *Bifidobacteriu*m and *Faecalibaculum* were most dominant during the pregnancy and lactation periods in MCD (Supplementary Fig. S2).

During the pups’ subsequent development, the PHFD uniquely harbored *Parabacteroides*, *Desulfovibrionaceae*, and *Bacteroides* during the 3^rd^, 4^th^, and 5^th^ weeks respectively. In contrast, the PMCD expressed a RA of beneficial bacteria such as *Muribaculaceae*, *Faecalibaculum*, *Rikenellaceae_RC9_gut group*, and *Bifidobacterium* during the same time points of development.

### 3.4 Impact of breastmilk on GM-specific vertical transfer and its stability

It is obvious that the most relatively abundant phylum in the GM of pups across their development stages is *Firmicutes* and *Bacteroidota* (Fig. 2). To better understand the impact of HFD on the vertical transfer of GM during offspring development, we assessed the RA of ASVs annotated to the class *Gammaproteobacteria* which are mostly pathogenic and prevalent in the environment and the order *Lactobacillales* which are prominent during the lactation stages. This study revealed high RA of *Lactobacillus* consistently transferred from dams to offspring during the lactation period in both maternal diets. In contrast, offspring from the MHFD group displayed a significant reduction in *Lactobacillus* RA after weaning in the 3^rd^ week, where the decline in *Lactobacillus* was accompanied by a gradual increase of *Enterococcus* and *Streptococcus* in the PHFD development time-points. However, the PMCD group contained GM-specific genera such as *Weissella*, while *Aerococcus* and *Streptococcus* are present in both pups (Fig. 4A).

**Figure 4:**
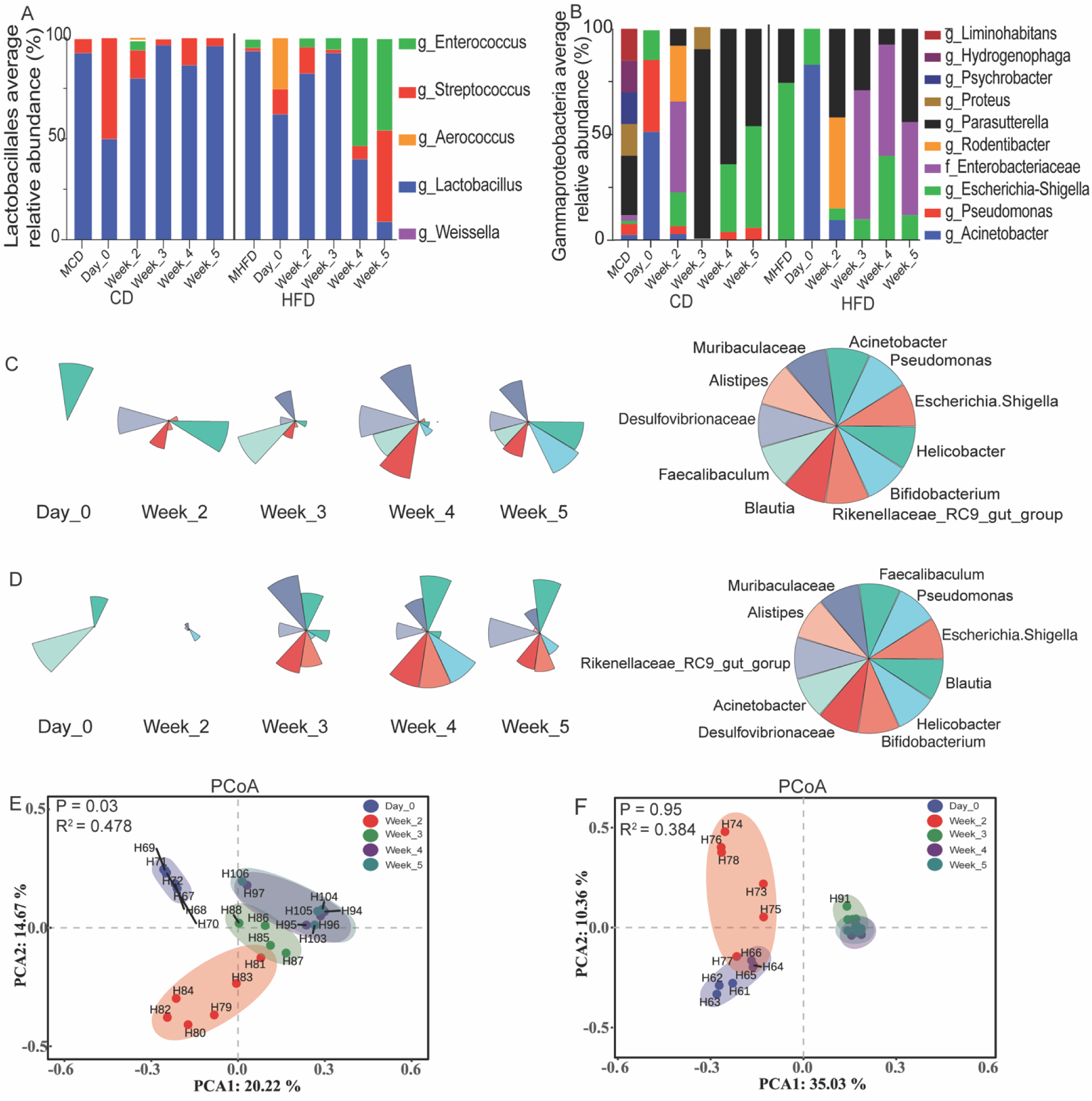
The impact of breastmilk in GM-specific vertical transfer to pups. Bar chart depicting the average proportion of taxa annotated to *Lactobacillales* (A) and *Gammaproteobacteria* (B) in the samples respectively. Figure keys were located on the right side of each chart. Pups’ fecal samples were grouped by GM and development stages. (B) PHFD genus contribution in the development process. (C) PMCD genus contribution in the development process. The corresponding range of each timepoint chart is represented on the right side of the group as pie chart. Data was calculated from ASVs annotation of the pups RA in the groups. Principal coordinate Analysis (PCA) among five pups’ development stages for PHFD (E) and PMCD (F) respectively. The percentage variance was indicated in the X and Y axis as PCA1 and PCA2, the significant difference was depicted as P<0.001. The colours represents the development stages at each time points.

During lactation, MCD fecal ASVs annotated to the class *Gammaproteobacteria* have several dominant genera such as *Parasutterella*, *Proteus*, *Psychrobacter*, *Hydrogenophaga*, and *Liminohabitans* which are not relatively expressed prior to lactation. Similarly, MHFD harbors only two genera of the class such as *Escherichia-Shigella* and *Parasutterella*, although only *Escherichia-Shigella* was found in the fecal ASVs of their pups prior to lactation. As the pups’ GM continued to develop, PMCD expressed a higher RA of *Parasutterella* and a lower abundance of *Escherichia-Shigella* a member of the *Enterobacteriaceae* family compared to PHFD that expressed a higher abundance of *Enterobacteriaceae* and one of its genus *Escherichia-Shigella*. Notably, PHFD has a lower abundance of *Parasutterella* as the GM continued to develop (Fig. 4B). Although the presence of *Escherichia-Shigella* was observed in all the pups from different dams, its relative abundance was low during the peak periods of lactation.

The transfer of specific GM from dams to their offspring varies according to diet. Both groups of pups exhibited a high RA of *Acinetobacter* prior to lactation. However, the pups from dams on a HFD displayed a notable presence of pathogenic bacteria, including *Helicobacter* and hydrogen sulfide-utilizing bacteria from the *Desulfovibrionaceae* family. In contrast, the pups from dams on a CD did not show significant abundance of these pathogenic bacteria or related phenotypes (Fig. 4A).

Instead, by the 3^rd^ week of GM development, PMCD exhibited a high RA of beneficial bacteria, specifically *Faecalibaculum*, *Muribaculaceae*, and *Bifidobacteria*. In comparison, the PHFD group showed only a moderate abundance of *Muribaculaceae* (Fig. 4B). As GM development progressed, the RA of *Faecalibaculum* continued to increase in the PMCD group, whereas its abundance decreased in the PHFD group. Conversely, in the PHFD group, the RA of *Muribaculaceae* progressively increased, while it diminished in the PMCD group.

As the pups matured into young adults, the GM of the PHFD group became compromised, characterized by a high RA of *Helicobacter*, a stark contrast to the gut microbiota observed in the PMCD group (Fig. 4A and 4B). This indicates that the dietary differences during lactation significantly influence the microbial composition and health of the offspring.

The PCA within the developmental stages of PHFD revealed variation in the microbial taxa at each timepoint, except for the 4^th^ and 5^th^ weeks, which showed early stability of the microbial taxa (Fig. 4E). Conversely, the microbial taxa of PMCD accounted for little or no variation, thus expressing early stability immediately after their dams’ weaning (Fig. 4F).

Post-hoc Bray-Curtis analysis further revealed the similarity of microbial taxa within each pup development stage. The PHFD results showed a dendrogram cluster with a higher rank of dissimilarity across day0 until after weaning during the development stages compared to PMCD group that exhibited similar microbial taxa during the 3^rd^, 4^th^, and 5^th^ weeks, showing low dissimilarity (Supplementary Fig. S3).

### 3.5 HFD induce a weaker correlation of specific microbial communities through association analysis

The vertical transfer of GM from dams to their pups, mediated by breast milk, revealed distinct differences in species assembly and community composition, suggesting specific ecological functions influenced by dietary variations. In the case of mothers on HFD and their pups, there was a weighted positive correlation with microbial taxa influenced by the HFD, characterized by a composition that included pathogenic bacteria centralized in modules M2 and M5. Conversely, other modules (M1, M3, and M4) were spatially decentralized, exhibiting negative correlations or complete disintegration from the microbial community. Notably, beneficial genera such as *Faecalibaculum*, *Muribaculaceae*, and *Lactobacillus* displayed negative correlations, indicating a lack of coordinated function (Fig. 5A).

**Figure 5:**
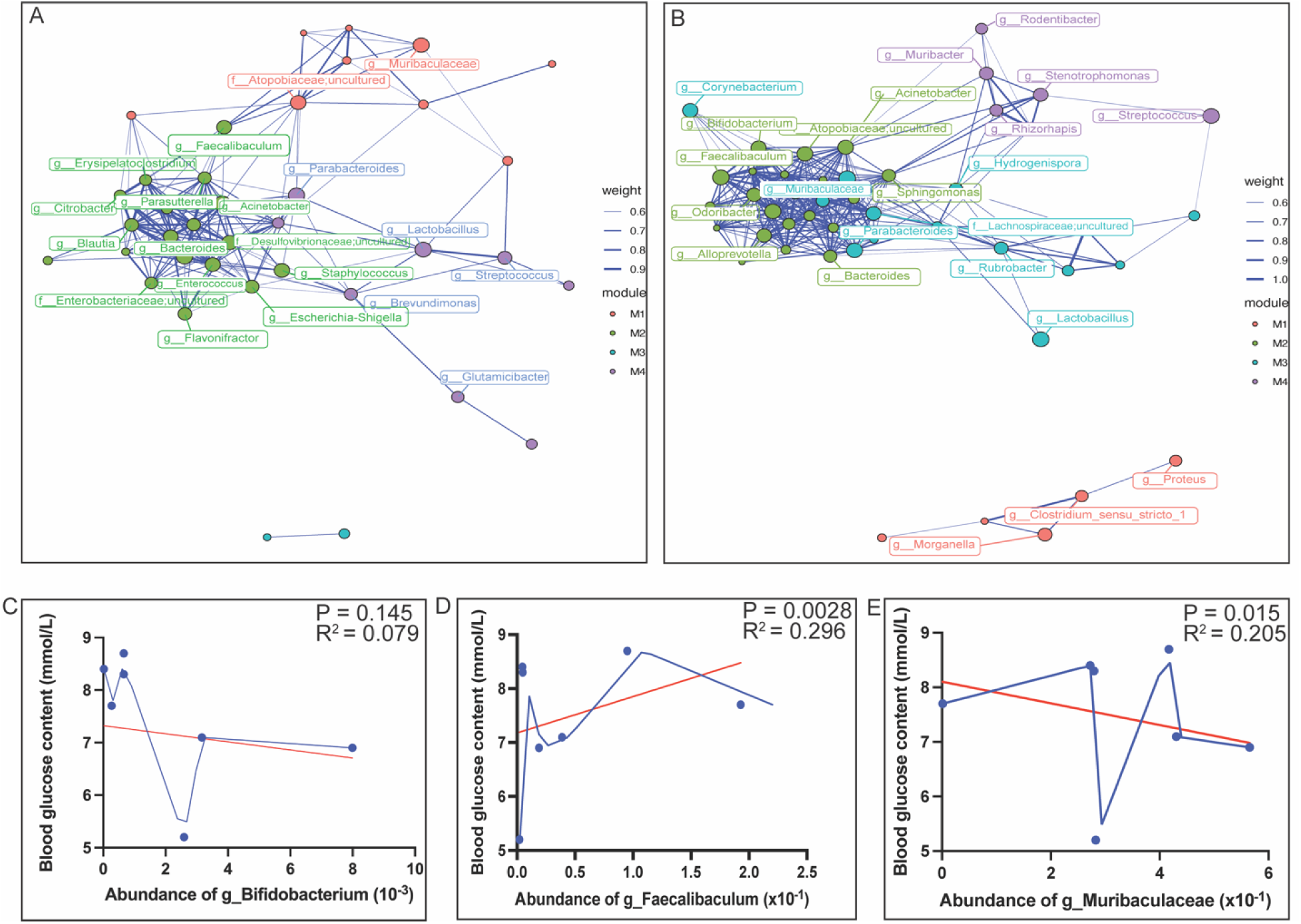
Microbial network analysis based on breastfeeding dams and lactating pups’ GM relationship. (A) igraph microbial network analysis showing the composition of the entire community, difference in assembly of species and their ecological function for HFD and (B) for CD. The nodes represent microbial taxa connected by edges of different thickness to indicate the correlation between different genera that are positive or negative. the distance between the nodes indicates the proximity of centrality and the number of times the node serves as a bridge between other nodes signify the importance of that microbial taxa. This spearman analysis was depicted as P<0.05, and 0.6 weighted edges indicate negative correlation while 0.9 or 1.0 indicates a positive correlation. Fit spline analysis showing the RA of some selected bacteria correlation with the blood glucose content of the mice (C) *Bifidobacteria*, (D) *Faecalibaculum*, (E) *Muribaculaceae*. P<0.05 was used to determine significant difference in this analysis.

In contrast, the mothers on a CD and their pups demonstrated a weighted positive correlation among beneficial microbial taxa, with the shortest proximal centralization observed in modules M2 and M3. These modules suggest a related coordinated function among the GM. However, other modules (M4 and M1) exhibited negative correlations with the centralized taxa and their network composition (Fig. 5B). The taxa in these modules indicate unique microbial functions distinct from the overall community.

Medically significant genera such as *Faecalibaculum*, *Bifidobacterium*, and *Muribaculaceae* showed positive correlations and connections with multiple taxa, facilitating a synergistic function influenced by the CD.

The fit spline analysis further suggested the impact of *Bifidobacterium*, *Faecalibaculum*, and *Muribaculaceae* on blood glucose levels. While the RA of *Bifidobacterium* revealed an insignificant relationship with maternal blood glucose levels (Fig. 5C), the abundances of *Faecalibaculum* and *Muribaculaceae* demonstrated moderate and significant relationships with blood glucose levels (Fig. 5D and 5E). Overall, these results highlight that while *Bifidobacterium* does not significantly influence blood glucose levels, both *Faecalibaculum* and *Muribaculaceae* exhibit meaningful correlations that glucose metabolism.

## 4.0 Discussion

The consumption of a HFD has been shown to negatively impact gut microbiota and exacerbate the progression of various chronic diseases [21]. Particularly concerning is the effect of HFD consumption by lactating mothers, which can induce hypertension in young offspring and reduce the RA of beneficial bacteria in infant GM [22,23]. This study observed that breastmilk produced by lactating mothers can significantly influence the developmental trajectory of their offspring through alterations in GM.

We found that the GM development in pups from dams on a HFD was notably different from that in pups from dams on a CD. This significant alteration in GM development was attributed to vertical microbiota transfer via breastmilk. Breastmilk as an initial diet is a crucial factor in the development of GM from infancy through to old age [24]. Our study revealed how breastmilk precede other vertical means of GM transfer to offspring, especially during pregnancy.

This highlights how offspring acquired GM can make them susceptible to GM-related changes that may occur in future such as delay in the establishment of a stable GM and potential risk of becoming diseased.

The microbial dysbiosis observed in pups from MHFD is closely linked to the dietary choices of the dams. This negative impact not only compromises the gut’s environmental conditions but also provides pathogenic bacteria with a selective advantage over beneficial bacteria. The proliferation of these pathogenic bacteria can predispose offspring to metabolic disorders and an unhealthy gut microbiome. Consistent with our findings, previous research has indicated that HFD can lead to GM-related issues, including social dysfunction and impaired synaptic plasticity [25]. Furthermore, the ASVs of pups annotated at birth and during lactation revealed significant differences in microbial diversity, highlighting the crucial role of breastmilk in the transfer of microbes from dams to their offspring. Although diet significantly affects the RA of phyla in pups, specific patterns emerged. For instance, *Firmicutes* and *Bacteroidota* exhibited higher RA in pups from the MCD group compared to those from the MHFD group, which demonstrated lower RA. This reduction in RA may be attributed to the competitive advantage of beneficial GM that thrive in nutrient-rich environments or possess superior clumping abilities. Consequently, we propose that the GM development in offspring of dams on a healthy diet is less susceptible to pathogenic and opportunistic bacterial genera. Supporting this perspective, Davis et al. (2022) reported that breastmilk substantially impacts both the taxonomic composition and functional capacity of the resident microbiome [26].

Our in-depth analysis focused on the maintenance of the average RA of *Lactobacillales* and *Gammaproteobacteria* throughout the developmental stages of pups, following vertical microbiota transfer via breastmilk from their dams. Notably, pups from the MCD group sustained their average RA of *Lactobacillus* throughout the entire development period. In contrast, pups from the MHFD group maintained their average RA only until weaning occurred. After weaning, we observed a gradual increase in the average RA of pathogenic bacteria such as *Enterococcus* in MHFD pups, which eventually displaced *Lactobacillus* as they developed into young adults. Similarly, the higher average RA of *Parasutterella* during the lactation period indicates effective vertical transfer of gut microbiota from dams to their offspring. However, the lower abundance of *Escherichia-Shigella* and the higher abundance of *Parasutterella* in pups from the MCD group suggest that non-lipogenic dams may positively influence the transfer of beneficial organisms during lactation. In contrast, the prevalence of *Escherichia-Shigella* and its family, *Enterobacteriaceae*, was minimized only during the peak of lactation in lipogenic dams.

Thus, we propose that breastmilk not only facilitates vertical microbial transfer to offspring but also possesses a unique protective mechanism that selectively maintains the colonization of beneficial GM, even under adverse conditions such as HFDs. This aligns with findings from previous studies, which indicate that breastmilk contains lactoferrin an essential factor influencing gut microbiome development, providing nourishment and immune protection while shaping GM [27,28]. Additionally, breastmilk is rich in prebiotics and immunological substances that promote the establishment of a favorable microbiome and dictate colonization patterns in infants [29,30]. It is noteworthy that lipogenic dams delay the GM stability of their offspring during their transition to young adulthood. In contrast, offspring from dams with a healthy diet achieve GM stability earlier, indicating a more normal and healthier microbiota development. These findings underscore the impact of lipogenic dams in promoting the growth of pathogenic microbial taxa, which can lead to microbial dysbiosis in their offspring. The gut microbial dysbiosis observed in pups from HFD dams can be attributed to various pathogenic bacterial genera, including *Escherichia-Shigella*, *Desulfovibrionaceae*, *Parabacteroides*, and *Bacteroides*. The persistence of these pathogenic bacteria throughout the development of MHFD pups may disrupt normal gut function by releasing inhibitory enzymes, competing for essential nutrients, or altering the gut environment necessary for the survival of beneficial organisms. This disruption negatively impacts the developmental processes of the pups. Although breastfeeding, particularly from lipogenic dams, has been reported to significantly influence the GM development of offspring in early life [31]. We identified dominant bacterial genera such as *Muribaculaceae*, *Faecalibaculum*, as potential probiotics that may be associated with glucose metabolism and contribute to gut homeostasis in juveniles. These genera may also play a crucial role in producing short-chain fatty acids, increasing sulfate production, and participate in amination and carboxylation which are beneficial for gut health. Consequently, researchers have recognized GM as a critical factor influencing the onset and progression of obesity, particularly concerning diet and host genetics [32,33]. The interplay between maternal diet and microbial transfer is critical in shaping the health trajectory of the offspring, highlighting the importance of a balanced gut microbiome in early life for proper growth and metabolic health.

## 5.0 Conclusion

The transfer of gut microbiota from parent to offspring is a crucial process that significantly influences health outcomes. This study highlights the essential role of breastmilk in facilitating this transfer between dams and their offspring. Our findings indicate that maternal diet plays a selective role in determining the composition of GM, whether pathogenic, beneficial, or opportunistic passed on to the young. Specifically, we discovered that lipogenic dams, through breastfeeding, can alter the GM development of their offspring, potentially increasing their susceptibility to various diseases as they mature into young adults. Interestingly, despite the impact of the maternal diet, breastmilk was effective in sustaining a healthy GM during the lactation period. This underscores the importance of breastmilk in promoting gut health, regardless of the dietary choices made by the dams.

## 6.0 Data availability statement

The data presented in this study are deposited in the Sequence Read Archive (https://www.ncbi.nlm.nih.gov/sra), under accession number PRJNA1142233 (http://www.ncbi.nlm.nih.gov/bioproject/1142233).

## 7.0 Ethics statement

The animal study was reviewed and approved by The IRB of School of Basic Medical Sciences of Central South University (No: CSU-2024-0315).

## 8.0 Authors contribution

ZY conceived the study. AU, and SL, JL and JH performed the experiments and analyzed the data. AU, JL and ZY wrote the manuscript, SL and HC participated in revision and language polishing. All authors contributed to the final version and approved the submission.

## 10.0 Acknowledgements

We thank Lei ShiBo for the technical support provided during the experiment. We also thank all the participants in this research.

## 12.0 Supplementary materials

**Supplementary figure S1:**
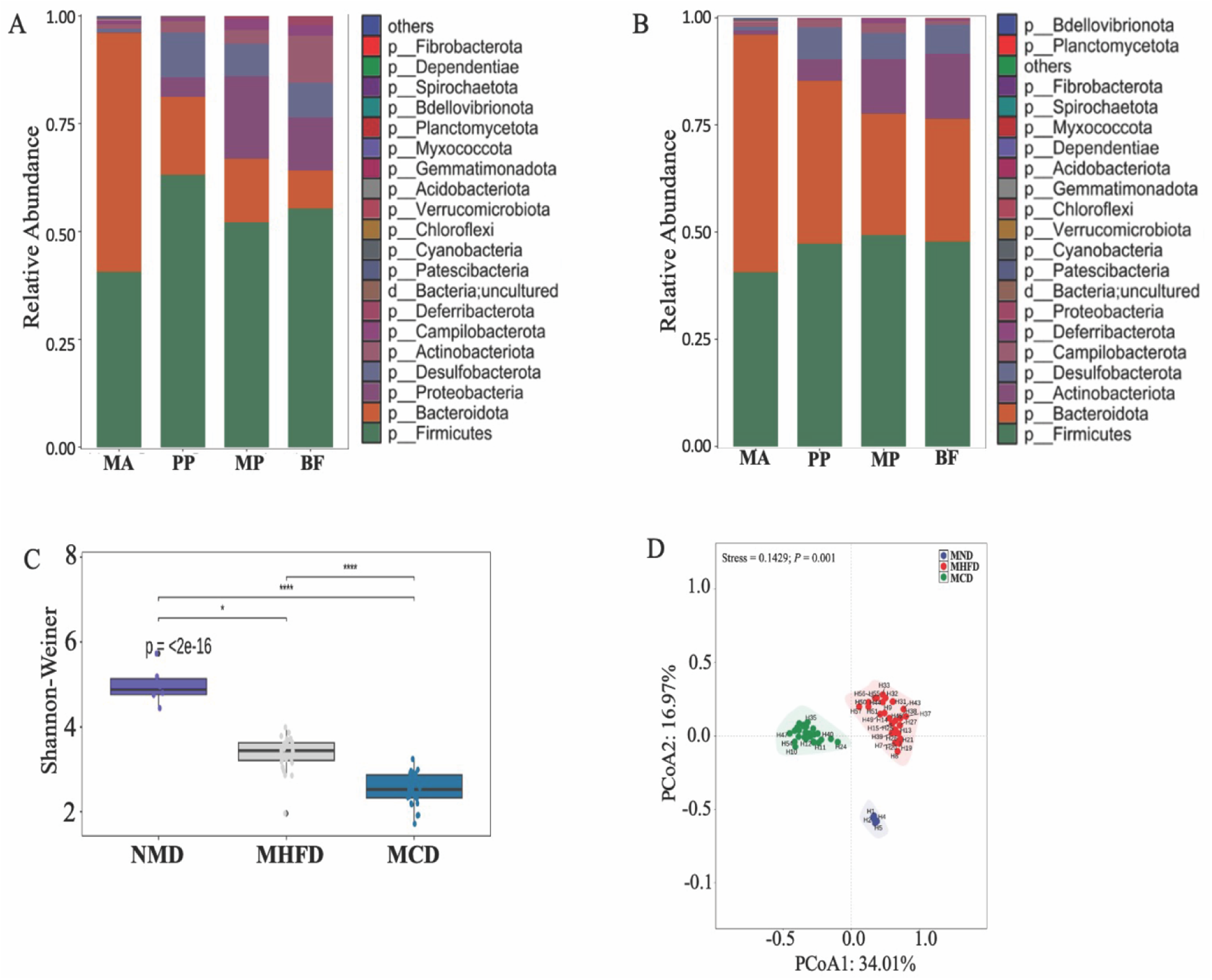
Consumption of HFD altered the diversity and composition of the maternal gut microbiota. (A and B) indicated changes in the relative abundance of phyla during the experimental periods. (C) The Shannon-weiner index with significant changes in the experimental groups (Normal, high-fat, and control diets). (D) Analysis of PCoA among the three groups. Different colours represented each treatment. Using multiple t-test comparison, **** indicates P < 0.0001, and * indicates P < 0.05. Dams’ adaptation (MA), Pre-pregnancy (PP), Maternal Pregnancy (MP), Breastfeeding (BF) and Normal maternal diet (NMD).

**Supplementary figure S2:**
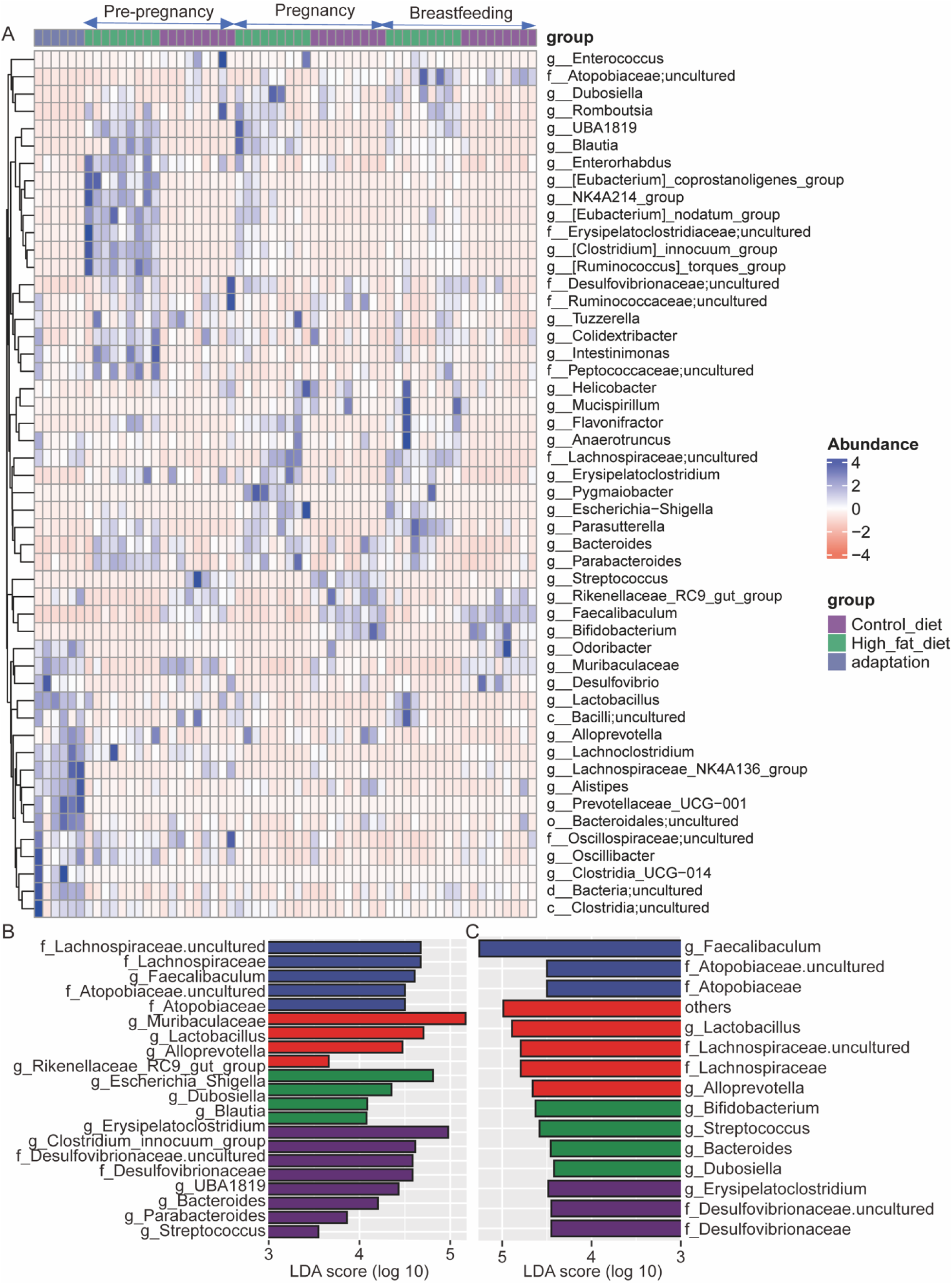
Maternal genera with significant differences in abundance level. (A) cluster heat map showing the abundance level of genera from different and similar taxonomic group. Different colours indicate significant changes in the relative abundance of different groups. The blue colours represent highly significant change in corresponding genera. (B and C) A histogram with linear discriminant analysis (LDA) scores in four groups for MHFD and PMCD respectively. Taxa highlighted in different colours indicate overrepresentation in the corresponding groups. The threshold of significance is set at 0.05. The threshold of the LDA score is set at 4.0.

**Supplementary figure S3:**
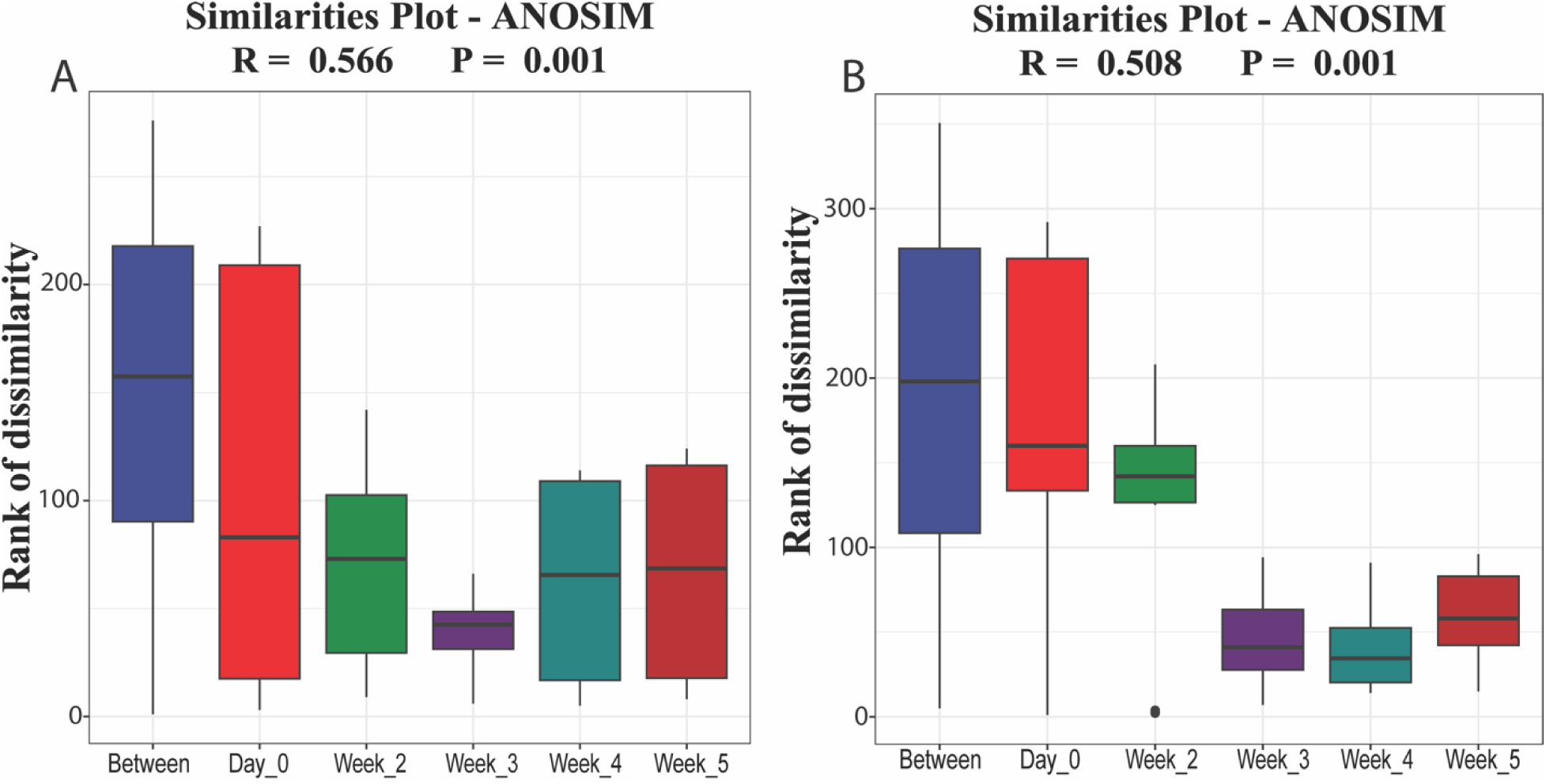
Pups’ fecal microbial composition was like other groups during GM development. An ANOSIM-similarity plot showed the rank of dissimilarity of gut microbiota composition between the four groups during developmental periods in PHFD (A and PMCD B) respectively. Dissimilarity between developmental periods was highlighted in different colours indicating the rank of dissimilarity in the groups. The threshold of significance was set at P value (0.001).

